# C-bouton components on rat extensor digitorum longus motoneurons are resistant to chronic functional overload

**DOI:** 10.1101/2021.02.05.429939

**Authors:** Roger W.P. Kissane, Arash Ghaffari-Rafi, Peter G. Tickle, Samit Chakrabarty, Stuart Egginton, Robert M. Brownstone, Calvin C. Smith

## Abstract

Mammalian motor systems adapt to the demands of their environment. For example, muscle fibre types change in response to increased load or endurance demands. However, for adaptations to be effective, motoneurons must adapt such that their properties match those of the innervated muscle fibres. We used a rat model of chronic functional overload to assess adaptations to both motoneuron size and a key modulatory synapse responsible for amplification of motor output, C-boutons. Overload of Extensor Digitorum Longus (EDL) muscles was induced by removal of their synergists, Tibialis Anterior (TA) muscles. Following 21 days survival, EDL muscles showed an increase in fatigue resistance and a decrease in force output, indicating a shift to a slower phenotype. These changes were reflected by a decrease in motoneuron size. However, C-bouton complexes remained largely unaffected by overload. The C-boutons themselves, quantified by expression of vesicular acetylcholine transporter, were similar in size and density in the control and overload conditions. Expression of the post-synaptic voltage-gated potassium channel (K_V_2.1) was also unchanged. Small conductance calcium activated potassium channels (SK3) were expressed in most EDL motoneurons, despite this being an almost exclusively fast motor pool. Overload induced a decrease in the proportion of SK3^+^ cells, however there was no change in density or size of clusters. We propose that reductions in motoneuron size may promote early recruitment of EDL motoneurons, but that C-bouton plasticity is not necessary to increase the force output required in response to muscle overload.

## Introduction

Evolution of the mammalian motor system affords many morphologically and functionally different animals to thrive in diverse environmental conditions (Brownstone, 2020). Yet the constantly changing environment – intrinsic and extrinsic – requires that an organism’s motor systems are capable of functional adaptations.

The muscular system itself is responsive to chronic changes in functional demand. For example, aerobic exercise training improves endurance capacity through coordinated increases in muscle activation patterns, which drives targeted, functional adaptation of skeletal muscle, altering metabolic signalling and expanding the vascular bed (Jensen et al., 2004). Additionally, chronic increases in loading drive muscle hypertrophy and increase force capacity (Mitchell et al., 2012). Moreover, constitutive fibre types, usually classified as slow oxidative (S-Type I), fast fatigue resistant (FR-Type IIA) and fast fatigable (FF-Type IIB/X) can undergo adaptive changes that alter muscle functional properties. For example, endurance training increases the proportion of Type I and IIA oxidative fibres resulting in improved fatigue resistance (Green et al., 1983).

Centrally, motoneurons innervating skeletal muscle also adapt to changing demands. These adaptations include changes in electrophysiological properties: for example, endurance training in rats induces increases in the motoneuron medium afterhyperpolarisation (mAHP) amplitude, consistent with a shift to a more fatigue-resistant phenotype (Gardiner et al., 2006). But there are also changes in organisation of synaptic inputs to motoneurons and in expression of postsynaptic membrane proteins, both of which reflect changes in neuromuscular activity (Woodrow et al., 2013, Arbat◻Plana et al., 2017). Thus, adaptations in spinal cord physiology parallel changes in the muscular system.

Motoneurons receive neuromodulatory cholinergic synapses, termed C-boutons because of their association with post-synaptic subsurface cisternae (Conradi, 1969), that regulate the mAHP (Miles et al., 2007). These somatic/proximal dendritic synapses can amplify motor output in a task-specific manner. The system is comprised of V0_C_ premotor interneurons (Zagoraiou et al., 2009), C-bouton synapses, and clusters of several different post-synaptic membrane proteins (Witts et al., 2014). Recent work has suggested mechanisms for how some of these components may contribute to amplification of motor output during high force output tasks such as swimming (Romer et al., 2019, Soulard et al., 2020, Nascimento et al., 2020). Activation of type 2 muscarinic acetylcholine receptors (m2AChR) on spinal motoneurons results in a reduction in the mAHP amplitude and an increase in excitability as measured by the frequency/current (ƒ-I) relationship (Miles et al., 2007). The mAHP current is carried by small conductance, calcium activated potassium channels SK2 & SK3, which are differentially expressed in fast and slow motoneurons, endowing them with their respective mAHP characteristics (Deardorff et al., 2013). Also clustered opposite C-boutons are delayed rectifier, voltage gated potassium channels, K_V_2.1 (Muennich and Fyffe, 2004). The role of these channels is less well understood, but recent evidence suggests they may act as ‘molecular rheostats’, capable of maintaining firing during high synaptic drive or supressing firing to protect motoneurons from excitotoxicity (Romer et al., 2019). It has been shown that m2AChRs modulate K_V_2.1 channels by reducing action potential half-widths and increasing the inter-spike AHP, which aids recovery of Na^+^ channels during high synaptic drive; thus supporting high frequency firing (Nascimento et al., 2020). Furthermore, as in various brain regions (Park et al., 2006, Murakoshi et al., 1997), motoneuron K_V_2.1 channels are plastic, suggesting they may play a role in neuromuscular adaptation in health and disease (Romer et al., 2014). However, the functional significance of K_V_2.1 channels in behaviour has yet to be determined.

Several groups have studied plasticity of C-boutons in disease states. For example, Landoni et al. (2019) showed that C-bouton transmission initially compensates for progression of motor deficits during motoneuron loss in SOD1 ALS mice. Conversely, Konsolaki et al. (2020) have shown that C-bouton inactivation improves motor performace but not survival in SOD1 ALS mice. It is difficult, however, to separate mechanisms associated with disease or injury and chronic physiological overload of the neuromuscular system. Therefore, it is important to study how chronic changes in neuromuscular demand affect central components of the motor system, such as the motoneuron and its modulatory inputs.

Here, we asked whether such changes in muscular demand lead to corresponding adaptations in motoneurons and at C-bouton synapses. We used a model of chronic neuromuscular overload, as similar models have previously been shown to induce central and peripheral adaptations in motor units (Ianuzzo et al., 1976, Rosenblatt and Parry, 1992, Krutki et al., 2015, Chalmers et al., 1991). This involved extirpating the tibialis anterior (TA) muscle to increase loading of the remaining synergist extensor digitorum longus (EDL) muscle in adult rats for 21 days. We confirmed effectiveness of the overload stimulus by assessing changes in muscle physiology, showing a shift to a more fatigue-resistant phenotype. We then studied EDL motoneuron adaptations using retrograde tracers, and showed a corresponding reduction in cross sectional area. Although we hypothesised that overload would induce adaptations to C-bouton organisation that correlated with adaptations seen in muscle physiology, there were no measurable differences in sizes and densities of both pre-(C-boutons) and post-synaptic (K_V_2.1 & SK3) components, however there was a reduction in the proportion of SK3^+^ cells following overload. Our results suggest that in conjunction with a slower muscle phenotype following overload, there is a corresponding decrease in motoneuron size. We suggest that this central adaptation may compensate for increased functional demands by reducing motoneuron rheobase and increasing excitability. Furthermore, anatomical plasticity of the neuromodulatory C-bouton complex is not necessary to produce increased force output in this model of chronic functional overload.

## Methods and materials

### Ethical approval

All surgical and experimental protocols were approved by the University of Leeds Animal Welfare and Ethics Committee and conducted in accordance with United Kingdom (UK) Animals (Scientific Procedures) Act 1986 (ASPA). The investigators understand the ethical principles under which the journal operates and confirm this work complies with the journal animal ethics guidelines.

### Animals

Male Wistar rats, (N = 14; 283 ± 29 g) were housed under a 12:12 light-dark cycle in a temperature-controlled 21°C environment, with *ad libitum* access to food and water. Animals were randomly allocated to either control of overload conditions.

### Animal surgical procedures

All animal surgeries were completed by competent Home Office approved PIL holders, under aseptic conditions. Surgical anaesthesia was induced and maintained with isoflurane (5% and 2%, respectively, in 100% O_2;_ IsoFlo®, Zoetis UK Ltd, London, UK).

### Muscle overload

An incision was made two thirds up the length of the right TA, towards the lateral side of the muscle. The covering fascia was cleared exposing the TA muscle, enabling sectioning of the distal tendon above the retinaculum and as close as possible to the proximal insertion (Egginton et al., 2011). Upon releasing the tendons, the TA was bluntly dissected from the lateral tibia surface and removed, taking care not to damage the underlying EDL. Skin was closed with 5-0 Mersilk suture (Ethicon, Johnson & Johnson Medical Ltd, New Brunswick, NJ, USA). Animals received subcutaneous analgesia (0.015mg/kg, Vetagesic®, Ceva, Amersham, UK) and antibiotic (2.5mg/kg, Baytril®, Bayer, Reading, UK) for two days post-surgery.

Removal of the TA muscle increases the load burden on its synergist, the EDL. We thus use the term “overload” for this condition. The overload period lasted 21 days.

### Motoneuron tracing

Retrograde fluorescent tracers were injected into the EDL five days prior to terminal experiments. Tracers were injected into the medial and lateral compartments of the EDL for separate assessment of motoneurons innervating these compartments (data not included). 1μL of 1.5% 647nm CTβ Alexa Fluor™ Conjugate (Invitrogen, Carlsbad, CA, USA) was injected into both medial and lateral compartments; 3μL of 1.5% Fast Blue (Polyscience, Inc., Warrington, PA, USA) was injected only into the medial compartment. Skin was closed using 5-0 Mersilk suture (Ethicon, Johnson & Johnson Medical Ltd, New Brunswick, NJ, USA). Animals received analgesic (0.015mg/kg, Vetagesic®, Ceva, Amersham, UK) and antibiotic (2.5mg/kg, Baytril®, Bayer, Reading, UK) subcutaneously for two days post-surgery.

### *In-situ* muscle fatigability

Anaesthesia was induced with isoflurane (4% in 100% O_2_) and maintained by constant infusion (30-35 mg kg^−1^ hr^−1^) of alfaxalone (Alfaxan: Jurox, Crawley, UK) via a catheter implanted into the external jugular vein. A tracheotomy was performed to facilitate spontaneous breathing. Blood pressure and heart rate were monitored in LabChart 8 (AD Instruments, UK) via a carotid artery catheter connected to a pressure transducer (AD Instruments, UK).

EDL twitch force was quantified using a lever arm transducer system (305B-LR; Aurora Scientific, Aurora, ON, Canada) and LabChart 8 (AD Instruments, UK). Unimpeded access to the EDL was enabled by dissection of surrounding fascia, the distal tendon was then cut and attached to the lever arm of the force transducer. The peroneal nerve was exposed and indirectly stimulated using bipolar stainless steel electrodes (Hudlicka et al., 1977), with muscle length and electrical current delivery optimised to generate maximal isometric twitch force. Fatigue resistance of the EDL was determined using a protocol (10 Hz electrical stimulation, 0.3ms pulse width) to elicit a series of isometric contractions over 3 minutes. A fatigue index (FI) was calculated as the ratio of end-stimulation tension to peak tension (FI = end-stimulation tension / peak tension), using the mean of 5 consecutive twitches

### Tissue preparation

Following successful *in situ* recordings, and remaining under anaesthesia, EDL were dissected and the muscle mid-belly was snap frozen in liquid nitrogen cooled isopentane for muscle capillary analysis. All frozen muscle tissue was stored at −80°C until cryo-sectioning. Next, animals were transcardially perfused with 0.1 M phosphate buffer and fixed with 4% paraformaldehyde. Spinal columns were removed immediately after perfusion and post-fixed in 4% PFA for 24 hours. Spinal cords were carefully dissected and cryoprotected in 30% sucrose at 4°C for 7 days. Next, the lumbar segments were isolated, frozen in OCT (Agar Scientific, Essex, UK) and stored at −20°C.

### Muscle processing

EDL muscles were cryo-sectioned (−20°C, 12μm), mounted on polylysine-coated slides (VWR International, Lutterworth, Leicestershire, UK), and stored at −20C until staining. Fibre boundaries were labelled with anti-laminin antibodies (Sigma-Aldrich, L9393) to identify the basement membrane. Capillaries were labelled by *Griffonia simplicifolia* lectin I (Vector Laboratories, FL-1101), an endothelial cell carbohydrate-binding protein. Photomicrographs were taken via a QImaging MicroPublisher 5.0 RTV camera (Teledyne QImaging, Surrey, BC, Canada) on a Nikon Eclipse E600 microscope (Nikon, Tokyo, Japan) at 20x magnification (field of view 440×330 μm) using a 2-second exposure time.

Indices for capillary-to-fibre ratio (C:F) and capillary density (CD) were derived from histological sections. These global indices describe gross changes in capillary supply, however they lack descriptive power of local capillary distribution. The local capillary supply is a critical determinant of functional capacity and of significant importance in the functional overload model which presents with a significant angiogenic response and fibre hypertrophy (Kissane et al., 2020, Tickle et al., 2020). Therefore, to investigate the influence of concurrent expansion of the capillary bed and fibre hypertrophy on muscle function, we mathematically modelled skeletal muscle oxygen transport kinetics (Al-Shammari et al., 2019). Briefly, capillary distributions were digitally derived from histological sections and used to model as a point source of O_2_ and estimations of tissue PO_2_ were predicted using a number of model assumptions: oxygen demand (15.7 ×10^−5^ ml O_2_.ml^−1^.s^−1^), myoglobin concentration (10.2 ×10^−3^ O_2_ ml^−1^), oxygen solubility (3.89 ×10^−5^ ml O_2_ ml^−1^.mmHg^−1^), myoglobin diffusivity (1.73 ×10^−7^ cm^2^ s^−1^) and capillary radius (1.8-2.5 ×10-4 cm; Al-Shammari et al., 2019).

We performed histological assessments of the fibre type distributions in both conditions, but tracer loading of the muscle reduced sample quality. These data were therefore excluded.

### Spinal cord immunohistochemistry

Spinal cord immunohistochemistry was performed as previously described (Smith et al., 2017). In brief, L3-L6 segments were sectioned at 50 μm on a cryostat (−20°C) and free-floating sections were collected and stored in PBS until staining. These were then washed in PBS (3 × 10 min) and incubated for 1-hour in blocking solution (0.2% Triton X-100, PBS, NaCl, and 10% normal donkey serum). The free-floating sections were then incubated for 48 hours in primary antibodies diluted in blocking solution, washed, and then incubated in secondary antibodies for 2 hours, also in blocking solution. Primary antibodies: goat anti-vesicular acetylcholine transporter (anti-VAChT, Millipore Cat# ABN100, RRID:AB_2630394, 1:1000), mouse anti-K_V_2.1 (UC Davis/NIH NeuroMab Facility Cat# 73-014, RRID:AB_10672253, 1:200), rabbit anti-SK3 (Millipore Cat# AB5350-200UL, RRID:AB_91797, 1:200). Secondary antibodies at 1:200: Alexa Fluor® 555 donkey anti-mouse (Thermo Fisher Scientific Cat# A-31570, RRID:AB_2536180), Alexa Fluor® 488 donkey anti-goat (Jackson ImmunoResearch Labs Cat# 705-546-147, RRID:AB_2340430), and Alexa Fluor® 555 donkey anti-rabbit 555nm (AB_2563181). Finally, sections were mounted on glass slides with Mowoil 4-88 (Carl Roth GmbH & Co. Kg, Karlsruhe, Germany).

### Confocal microscopy and quantitative analysis

Images were acquired with a Zeiss LSM 800 confocal microscope (Zeiss LSM 800 with Airyscan, RRID:SCR_015963), using a 40x oil immersion objective (1 AU aperture), and Zeiss ZEN Blue Edition software (ZEN Digital Imaging for Light Microscopy, RRID:SCR_013672). Motoneurons were identified by their location in the spinal cord ventral horn and presence of CTβ (647nm) or FB (405nm) staining. Z-stacks of 30 μm at 0.40 μm intervals were acquired through the centre of each neuron, identified by the nucleus. Motoneurons that did not have a visible nucleus or had significant membrane disruption were excluded from due to poor reconstruction quality.

Researchers were blinded to the conditions for all data analyses. 3-dimentional (3D) reconstructions of each motoneuron were rendered from the confocal image z-stacks, utilising IMARIS Software (IMARIS, RRID:SCR_007370). In the 3D isometric view, solid surfaces of the motoneuron soma with dendrites, C-boutons, and K_V_2.1 or SK3 were created via surface rendering and thresholding. CTβ or Fast Blue was used to model the motoneuron surface. A masking feature was then used to select K_V_2.1 or SK3 clusters contacting the motoneuron surface and/or proximal to the C-bouton. IMARIS was then used to generated volume and surface area data for each motoneuron, C-boutons, K_V_2.1 clusters and SK3 clusters. To determine the motoneuron cell size, cross-sectional area through the centre of the nucleus was calculated using Image J (ImageJ, RRID:SCR_003070).

Data were then exported to Excel (Microsoft Excel, RRID:SCR_016137). Since all alpha-motoneurons contain C-boutons and K_V_2.1, cells with no C-bouton or K_V_2.1 surfaces were removed (Deardorff et al., 2014b). C-bouton, SK3 and K_V_2.1 channel densities were normalised to the motoneuron surface area. For statistical analyses, the mean synaptic and channel density for each motoneuron was the observational unit (n) and the average density per animal was the experimental unit (N). Sample sizes were determined based on previous studies (Kissane et al., 2018, Romer et al., 2014). Shapiro-wilks tests were performed to determine normality of the data, followed by either unpaired t tests (normally distributed) or Mann-Whitney U tests (not normally distributed). Fisher’s exact test was used to compare the proportion of cells expressing SK3. Data are presented throughout as mean ± standard deviation. All analyses were performed using Python scripts (RRID: SCR_008394) in the Jupyter notebooks environment (RRID: SCR_013995, link to analysis notebook provided with publication).

## Results

### Chronic functional overload induces functional shift to slower EDL phenotype

Previous studies of EDL muscle overload by removal of the TA synergist have shown an anatomic and physiologic shift to a slower phenotype (Rosenblatt and Parry, 1993, Rosenblatt and Parry, 1992). We were not able to reliably analyse fibre type distribution in this study due to the loading of muscles with neuro-anatomical tracers. However, muscle weight was significantly greater in the overload condition, suggesting hypertrophy of EDL fibres (Control=0.06 ± 0.01g, N=5 vs Overload 0.09 ± 0.02g, N=7, p=0.001).

### Chronic functional overload improves fatigability in EDL muscles

In order to determine if there were physiological adaptations in EDL muscles following overload, we assessed muscle twitch force and fatigability using an *in vivo* anaesthetised preparation. Measurement of maximal isometric force showed that overloaded muscle had reduced twitch (227 ± 55 g/g, N=7 vs 296 ± 36 g/g, p=0.017, N=5, Fig. 1A, A^1^) and tetanic (Control= 1184± 189 g/g, N= 5 vs Overload= 864 ± 155 g/g, N=7, p=0.009, Fig. 1B, B^1^) force outputs compared to control. Correspondingly, the fatigue index was increased in the overload condition (0.64 ± 0.04) compared to control (0.50 ± 0.08, p=0.0003, Fig. 1C) indicating a shift to slower, more fatigue resistant fibres in the overload condition. Assessments of muscle capillary supply highlighted the potent angiogenic response to overload with a significant increase in capillary-to-fibre ratio (Control=1.67 ± 0.10, N=6; Overload=2.15 ± 0.35, N=5, p=0.01, unpaired t test, Fig. 1D). However, due to hypertrophy induced by overload, the global capillary density (Control=920 ± 122 mm^−2^, N=6; Overload=761 ± 196, N=5, p=0.13, unpaired t test, Fig. 1E) and modelled spatial tissue PO_2_ remained unchanged (Control=11.5 ± 1.8 mmHg N=6; Overload=9.1 ± 3.1 mmHg, N=5, p=0.12, unpaired t test, Fig. 1F-J). Thus, improved fatigue resistance is likely due to increased efficiency in aerobic metabolism.

**Figure 1.**
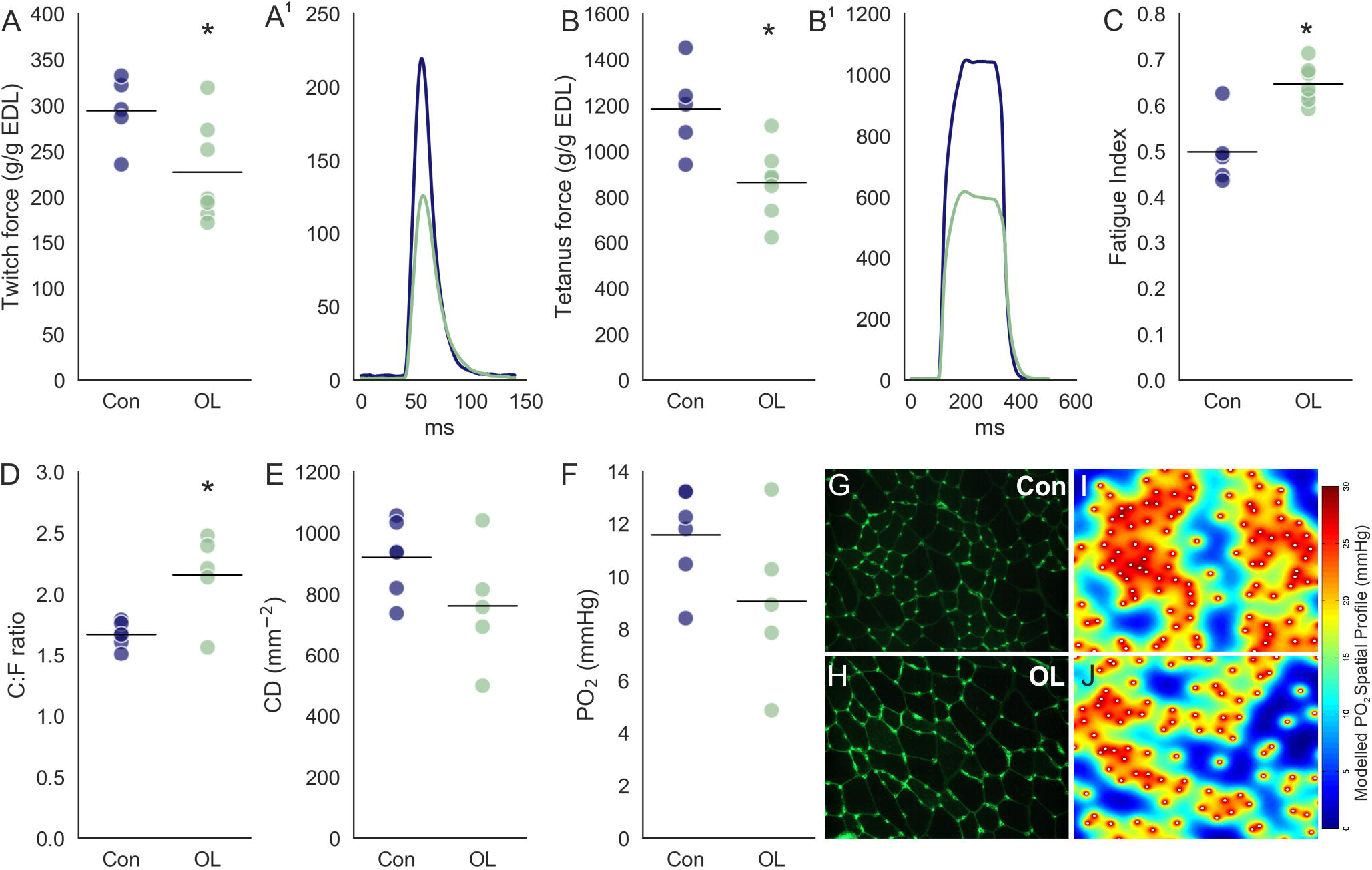
Effect of overload on EDL force production and fatigability. (A) Maximum twitch force in control and overload animals normalised to EDL weight (g/g). (A^1^) Representative normalised twitch traces from a control (blue) and overload (green) EDL. (B) Maximum tetanic force normalised to EDL weight (g/g). (B^1^) Representative peak normalised traces during tetanic stimulation in a control and overloaded EDL muscle (C) Fatigue index ratio (end-stimulation force/peak force). (D) Capillary-to-fibre ratio. (E) Capillary density (CD) per mm^−2^ of muscle fibre. (F) Modelled partial pressure of O_2_ (PO_2_). (G-H) Representative 20x light microscope images of lectin stained (capillaries) muscle fibres in control (G) and overload conditions (H). (I-J) Representative images of modelled PO_2_ spatial profile for control and overload conditions. Experimental units (animals, N) and statistical tests are as follows: A-C, control N= 7, overload N= 7, unpaired t test; D-F, control N= 6, overload N= 5, unpaired t test. * represents a statistically significant difference (p<0.05).

Overall, our physiological data confirm a shift to a slower, more fatigue resistant phenotype following overload.

### Chronic functional overload reduces EDL motoneuron soma cross section area

Motoneuron properties are matched to the muscle fibre types they innervate. Motoneurons innervating Type 1 muscle fibres are smaller than those innervating IIa, which are smaller than those innervating IIb/IIx (Burke, 1967). In the overload model, we found a shift in the distribution of motoneuron sizes towards smaller motoneurons, leading to an overall reduction in EDL motoneuron size following removal of TA (**Control**= 1749 ± 248, N= 6, n=285 vs **Overload**= 1372 ± 188, N=5, n=260, p= 0.021, unpaired t test, Fig. 2A-D^1^). This shift to smaller sized motoneurons corresponds to the shift in muscle phenotype.

**Figure 2.**
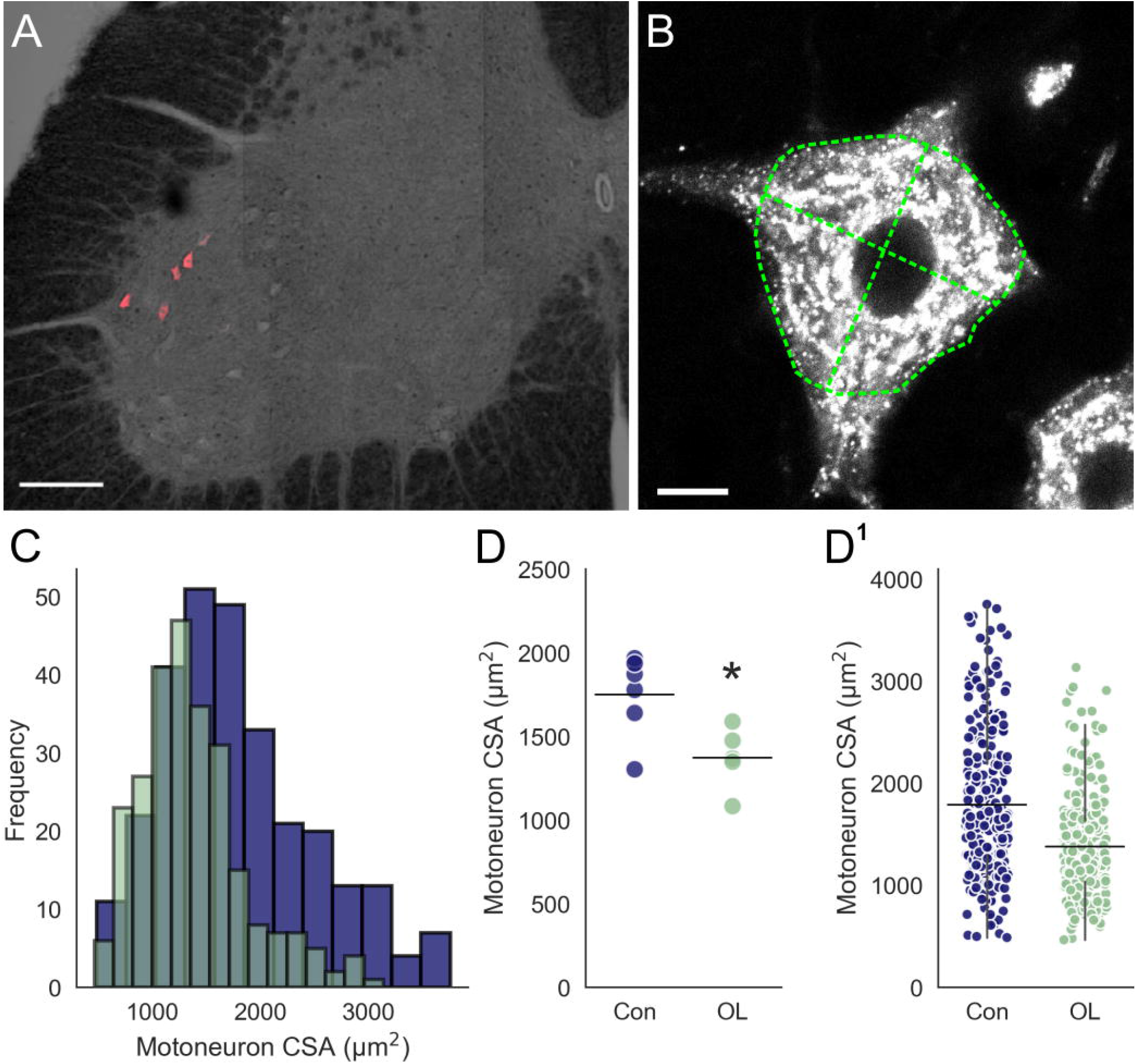
Changes in EDL motoneuron size following chronic overload. (A) 10x confocal tile scan image illustrating retrograde labelled EDL motoneurons (CTβ-647) on a transmitted light background to show section morphology. (B) Representative single optical slice, 60x confocal image through the centre of labelled motoneuron. Green dashed lines illustrate cell perimeter and cross sectional lines for measuring CSA. (C) Frequency distributions for CSA of control (blue) vs overloaded (green) motoneurons. (D) Strip-plots comparing mean motoneuron CSA in control and overload animals. (D^1^) Strip plot showing all motoneurons analysed from each group. Experimental units (animals, N) and statistical tests are as follows: D, control N= 6, overload N= 5, unpaired t test. * represents a statistically significant difference (p<0.05). Scale bar in A= 150 μm, a= 10 μm. Whiskers extend to 1.5 × SD of the mean.

### Overload has no effect on C-bouton innervation of EDL motoneurons

C-bouton synapses are terminals of the V0_C_ interneuron circuit responsible for task-specific amplification of motor output (Miles et al., 2007, Zagoraiou et al., 2009). Because the overload condition removed the contribution of the synergist, TA, to locomotion and necessitated increased force output from the EDL, we asked whether an increase in C-bouton synapses (Fig. 3A, A^1^) occurs in order to meet the increased demands placed on EDL motoneurons. We thus assessed the effect of overload on the density and size of C-bouton inputs.

**Figure 3.**
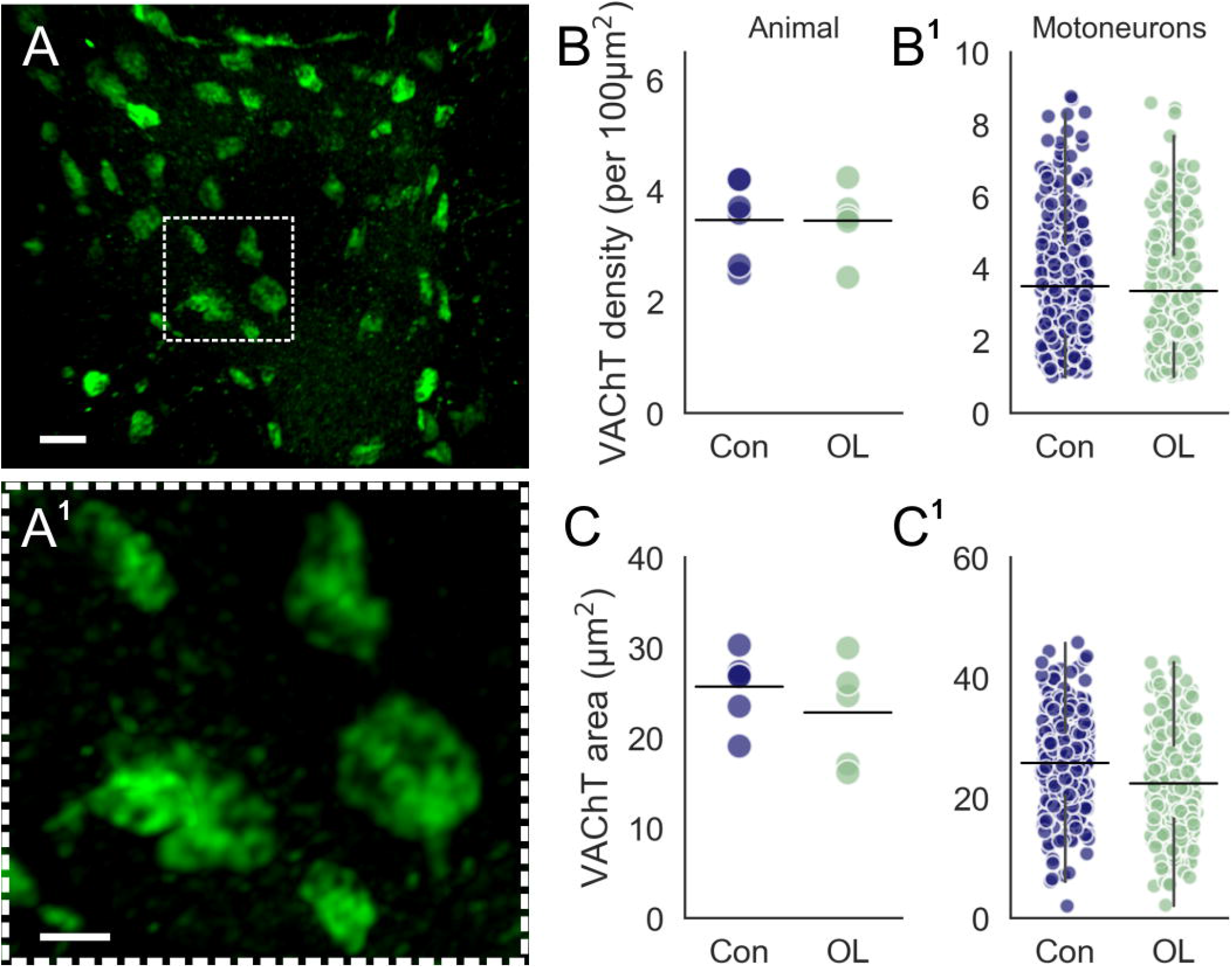
C-bouton innervation of EDL motoneurons following chronic overload. (A) Representative 3D projection of 60x z stack confocal image showing VAChT positive C-boutons on an EDL motoneuron. (A^1^) High magnification image of the demarcated region in A, showing C-boutons. (B) Strip-plot showing mean C-bouton density for control (blue) and overload (green) animals. (B^1^) Strip-plot of mean C-bouton density for individual motoneurons in control and overload conditions. (C) Strip-plot showing mean C-bouton area for control and overload animals. (C^1^) Strip-plots showing mean C-bouton area for individual control and overload motoneurons. Experimental units (animals, N) and statistical tests are as follows: B & C, control N= 6, overload N= 5, unpaired t tests. Scale bar=4 μm, a-c= 1 μm. Whiskers extend to 1.5 × SD of the mean.

There was no significant effect of overload on C-bouton density (Control= 3.4 ± 0.8 per 100 μm^2^, N=6, n=289 vs Overload= 3.3 ± 0.8 per 100 μm^2^, N=5, n=259, p=0.61, Fig. 3B, B^1^), or mean area (Control=25.7 ± 3.7 μm^2^, N=6, n=289 vs Overload =23.3 ± 6.9 μm^2^, N=5, n=259, p=0.48, Fig. 3C, C^1^). That is, there were no discernible changes to the presynaptic component of C-bouton inputs to EDL motoneurons.

### K_V_2.1 channel density and area are unaffected by overload

Although we saw no change in presynaptic C-bouton characteristics, changes in the post-synaptic protein complex could alter synapse function. K_V_2.1 channels are thought to be important for facilitating high frequency motoneuron firing and are recruited for C-bouton amplification of motor output (Nascimento et al., 2020). In addition to the large clusters found at the C-bouton, K_V_2.1 channels are found in smaller clusters distributed throughout the membrane (Fig. 4A-C^1^). We therefore assessed the influence of overload on expression of both C-bouton associated and unassociated K_V_2.1 channels by masking K_V_2.1 signal to the presynaptic VAChT signal.

**Figure 4.**
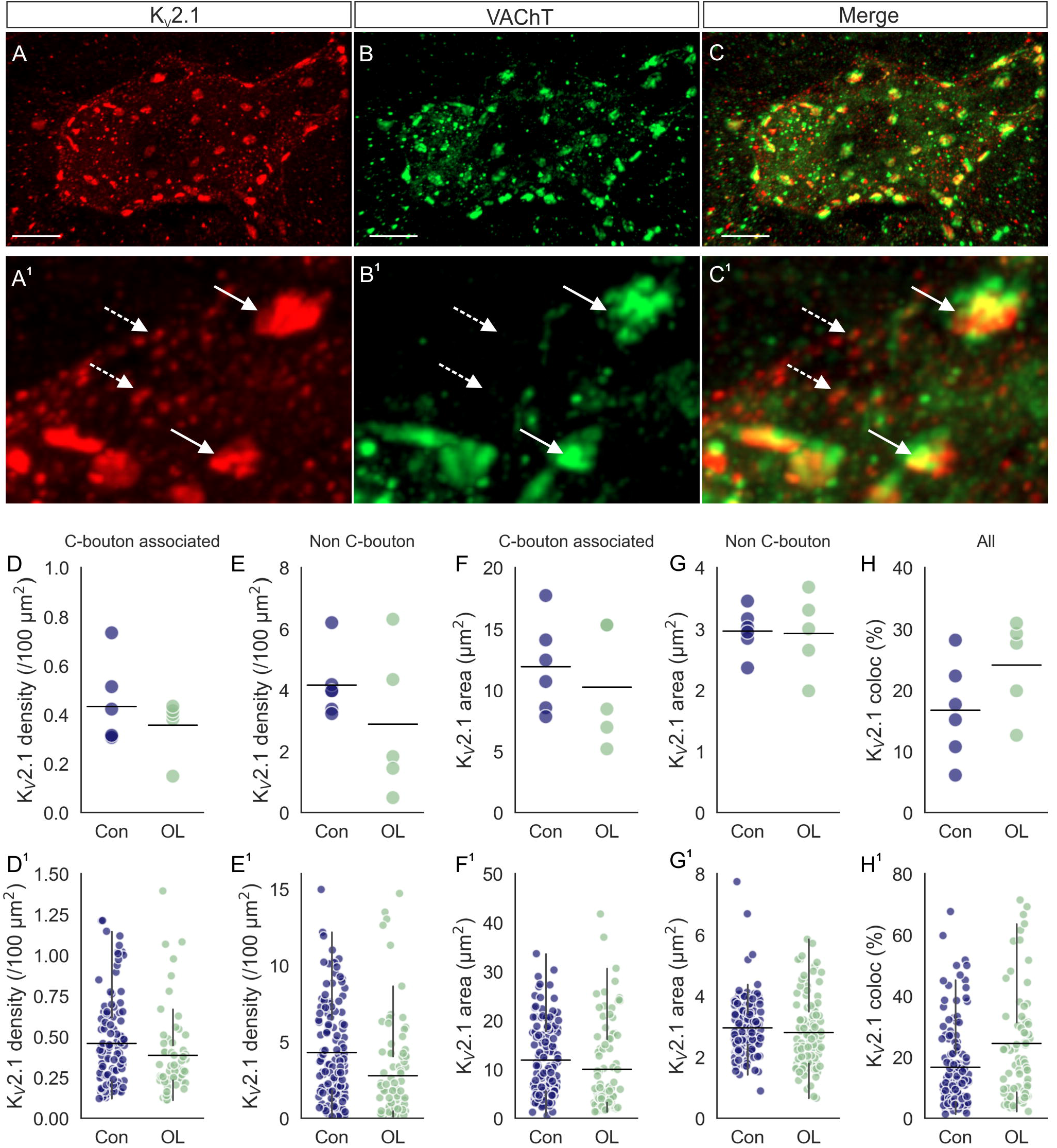
The effect of overload on K_V_2.1 expression. (A-C) Representative 3D projection of 60x z stack confocal image showing showing K_V_2.1 and VAChT fluorescence. (A^1^-C^1^) High magnification, 3D confocal z stack projections showing K_V_2.1 clusters in apposition with VAChT positive C-boutons (solid arrow), and those not (dashed arrow). (D) Strip-plot showing mean K_V_2.1 density in control (Con) and overload (OL) animals. (D^1^) Mean K_V_2.1 cluster density for individual motoneurons. (E-E^1^) As in D-D^1^ for density of C-bouton unassociated K_V_2.1 clusters. (F) Strip-plot showing mean surface area of K_V_2.1 clusters associated with C-boutons in control and overload animals. (F^1^) Mean area of C-bouton associated K_V_2.1 clusters for individual motoneurons. (G-G^1^) As in F-F^1^ for surface area of C-bouton unassociated K_V_2.1 clusters. (H) Percentage Kv2.1 co-localised to the C-bouton in control and overload conditions. (H^1^) As in H with all motoneuron means shown. Experimental units (animals, N) are as follows: D-H, control N= 6, overload N= 5. Statistical tests performed were as follows: D, F and H, unpaired t tests; E and G Mann WhitneyU tests. * represents a statistically significant difference (p<0.05). Scale bars in A-C= 10 μm, A^1^-C^1^= 1 μm. Whiskers extend to 1.5 × SD of the mean.

Chronic functional overload did not significantly alter the density of K_V_2.1 clusters either associated (Control=0.43 ± 0.17 per 100 μm^2^, N=6, n=135 vs Overload= 0.36 ± 0.12 per 100 μm^2^, N=5, n=85, p=0.41, t-test, Fig. 4D, D^1^), or unassociated (Control=4.2 ± 1.1 per 100 μm^2^, N=6, n=135 vs Overload= 2.9 ± 2.4 per 100 μm^2^, N=5, n=85, p=0.26, Fig. 4E, E^1^) with C-boutons. Similarly, there was no difference in mean surface area between control (associated: 12.4 ± 3.5 per 100 μm^2^; unassociated: 3.0 ± 0.4 per 100 μm^2^, N=6, n=135) and overload groups (associated: 10.1 ± 5.4 per 100 μm^2^, p=0.41, Fig. 4F, F^1^; unassociated: 3.0 ± 0.6 per 100 μm^2^, N=5, n=85, p=0.26, Fig. 4G, G^1^). The lack of change in density and surface area of K_V_2.1 for both associated and unassociated K_V_2.1 channels was reflected in the unchanged percentage of K_V_2.1 localised to the C-bouton following overload (Control=16.8 ± 16.8 per 100 μm^2^, N=6, n=140 vs Overload= 24.5 ± 21.2 per 100 μm^2^, N=5, n=79, p=0.15, Fig. 4H, H^1^).

### SK3 channels are expressed in most EDL motoneurons but are unaltered by chronic functional overload

In addition to K_V_2.1 channels, small conductance potassium channels (SK) are also clustered on the motoneuron post-synaptic membrane opposing C-boutons. SK2 and SK3 channels are responsible for the calcium-dependent potassium currents underlying the mAHP and therefore regulate motoneuron firing frequencies. Previous work has suggested that SK2 channels are expressed in all motoneurons, whereas in the specific pools studied (mainly tibial motoneurons), SK3 channels are selectively expresssed in slow motoneurons (Deardorff et al., 2013, Dukkipati et al., 2018). Although the EDL muscle is mainly comprised of fast fatiguable (FF) units, there are also fast fatigue resistant (FR) and a small proportion of type I slow (S) units (Kissane et al., 2018). We reasoned that, given the shift to a more fatigue resistant phenotype following chronic overload (Fig. 1), there would be a corresponding upregulation of SK3 expression in motoneurons.

Interestingly, we found that out of all motoneurons assessed (regardless of condition) 227 out of 285 (79%) expressed SK3 channel clusters (Fig. 5A-E). In control animals, 130 out of 141 cells expressed SK3 (92%), while in the overload condition 97 out of 144 motoneurons (67%, p=9.5e-06) had SK3 expression. The lower proportion of SK3 in the overload condition may indicate that SK3 is in fact downregulated following chronic overload, however in the neurons that expresed SK3, we found no difference between control and overload groups in either SK3 density (Control=0.4 ± 0.04 per 100 μm^2^, N=6, n=130 vs Overload=0.5 ± 0.3 per 100 μm^2^, N=5, n= 97, p=0.65) or area (Control=7.4 ± 2.7 μm^2^, N=6, n=130; Overload=7.0 ± 2.0 μm^2^, N=5, n=97, p=0.73). Thus, while fewer cells were SK3 positive in the overload condition, there was no difference in SK3 expression between the conditions for cells that were positive.

**Figure 5.**
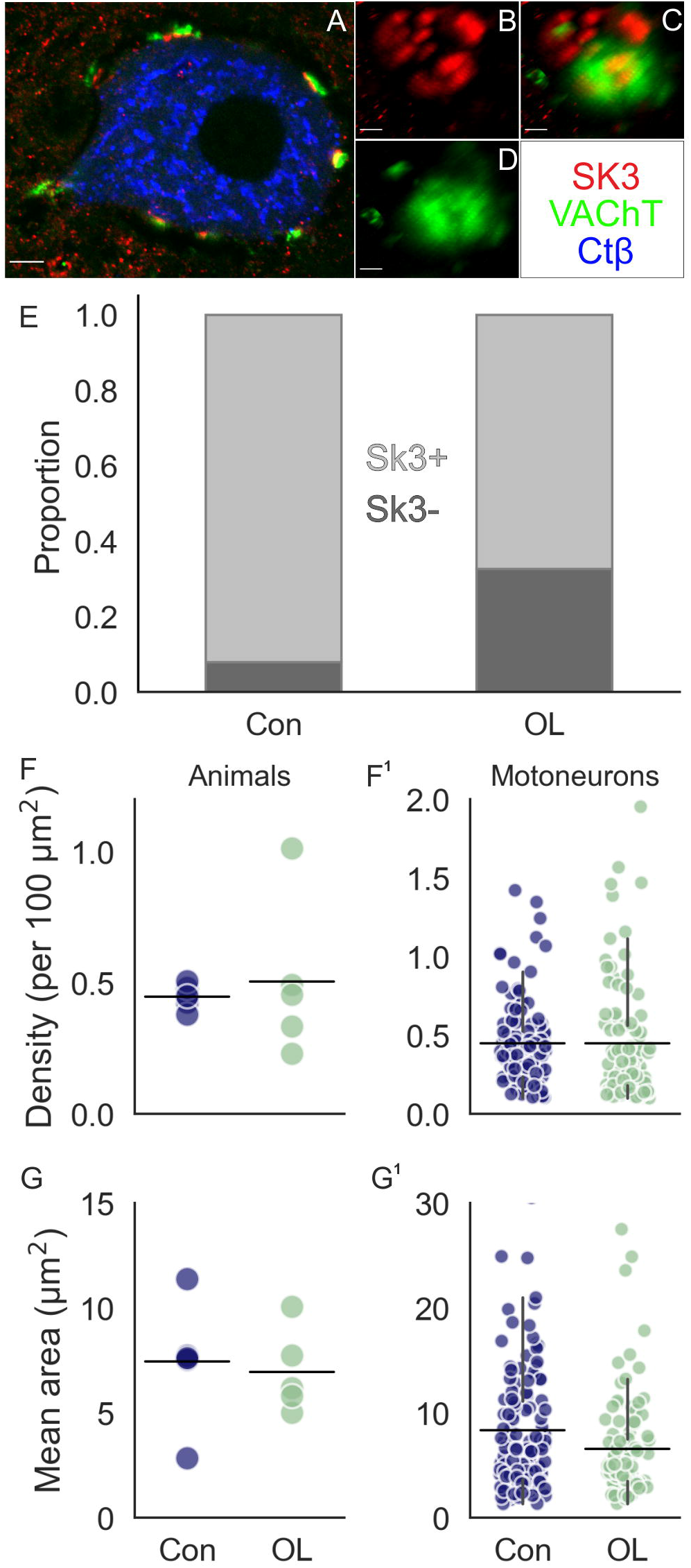
The effect of overload on SK3 expression on EDL motoneurons. (A) Representative 60x confocal image through the centre of a CTβ positive EDL motoneuron showing staining for SK3 (red) and VAChT (green). (B-D) high magnification image of SK3 cluster apposing VAChT positive C-bouton. (E) Proportion of all cells positive and negative for SK3 in control and overload conditions. (F) Strip-plot showing mean SK3 density for control (blue) and overload (green) animals. (F^1^) Strip-plot showing mean SK3 density for individual motoneurons. (G) Strip-plot showing mean SK3 area for control and overload animals. (G^1^) Strip-plot showing mean SK3 area for individual motoneurons. Experimental units (animals, N) and statistical tests are as follows: F-G, control N= 6, overload N= 5, unpaired t tests. Scale bars A =5 μm; B-D=0.5 μm.

In summary, we found that SK3 channels are not reliable binary markers for slow motoneurons in EDL and that there was a decrease in the proportion of SK3^+^ cells in the overload condition despite the shift to smaller cell sizes and decreased fatigability. Overall, though, we detected no significant effect of overload on presynaptic C-bouton terminals, or expression of either of the post synaptic proteins assessed (SK3 and K_V_2.1). This suggests that the C-bouton complex is resistent to the increased functional demands of the innervated muscle.

## Discussion

The results presented here demonstrate that chronic overload of the rat EDL muscle induces significant adadptations to muscle fibre capiliary innervationand contractile properties. Specifically, 21 days functional overload resulted in an increase in EDL capiliary-to-fibre ratio and fatigue resistance, parralelled by a decrease in twitch and tetanic force production. We reasoned that this shift to a slower phenotype in the overloaded EDL muscle may be reflected anatomically in their motoneuron characteristics. This was confirmed by a decrease in motoneuron soma cross-sectional area. We also hypothesised that a key spinal neuromodualtory input, C-boutons, would change so as to compensate for the increased neuromuscular demand during overload. However, while we did find a decrease in the proportion of SK3^+^ neurons, we were unable to detect significant adaptations in size or density of key components of the C-bouton complex (C-bouton synapse, SK3 and K_V_2.1 channels). Importantly, we also show that SK3, a suggested molecular marker for slow motoneurons (Dukkipati et al., 2018), was expressed in most EDL motoneurons, despite these being mainly fast type (Kissane et al., 2018).

### Increased EDL fatigue resistance following overload is paralleled by a reduction in motoneuron soma size

The overload induced shift to a more aerobic phenotype in EDL muscle was reflected in the central portion of the motor unit, by a shift to smaller sizes for motoneurons in this condition. Motoneuron size is inversely related to input resistance and positively correlates with rheobase (Henneman, 1957), meaning changes in motoneuron size may lead to altered recruitment thresholds across the motor pool. Previous electrophysiological assessments of overloaded rat medial gastrocnemius (MG) motoneurons demonstrated significant reductions in rheobase, associated with increased input resistance and a leftward shift in the frequency current (ƒ-I) relationship in FF motoneurons, indicating that less input was required to activate these motoneuorns (Krutki et al., 2015). Taken together, these changes in motoneuron properties would mean that the probability of recruitment and firing rate is increased for a given synaptic drive in the overload compared to the control condition. This would be a beneficial compensatory adaptation to the increased demands induced by the removal of TA muscle.

### C-bouton complex is largely unaltered by chronic functional overload

The C-bouton is a neuromodulatory synapse responsible for task specific amplification of motor output, and is anatomically characterised by a dense aggregation of proteins at the opposing post synaptic membrane (Conradi, 1969, Deardorff et al., 2014a). C-bouton modulation increases the ƒ-I slope of motoneurons through activation of m2AChRs and subsequent reductions in the amplitude and duration of the mAHP, mediated by SK channels (Miles et al., 2007). C-bouton SK2 channels are expressed in almost all motoneurons regardless of type, whereas previous work in extensor motoneurons shows that SK3 channels are preferentially found on S type motoneurons and thus are likely responsible for larger mAHP conductances (Deardorff et al., 2013, Dukkipati et al., 2018). Surprisingly, we found that most EDL motoneurons expressed large SK3 clusters to some degree. Thus, in the rat EDL motor pool, it would seem that SK3 expression cannot be reliably used as a binary molecular marker to identify slow motoneurons.

There was a decrease in the proportion of SK3^+^ motoneurons in the overload condition. Given prior evidence that SK3 channels are associated with longer AHPs and slower phenotypes (Deardorff et al., 2013), the “loss” of SK3 expression could be considered to indicate a shift to a faster phenotype – a finding in contrast to our other findings. However, considering that we did not detect a decrease in SK3 expression (cluster size or density) in the SK3-positive cells, it is possible that the motoneurons that presumably ‘lost’ SK3 clusters had lower expression levels before overload. Taken together, it is not clear whether the reduction in the number of SK3^+^ neurons had significant functional implications.

We found no effect of overload on the size or density of C-boutons, K_V_2.1 channels, or SK3 channels on EDL motoneurons, suggesting that both pre and post-synaptic components of the synaptic complex are largely unaffected by a stimulus which induced changes in muscle fibres and motoneuron size.

This was unexpected, especially for SK3, as the motoneuron mAHP has been shown to be increased following chronic overload (Krutki et al., 2015). It is possible that there were changes in other SK isoforms, but given the propensity of SK3 channels to be differentially expressed in motoneuron types, we focused on them.

There are several potential reasons why we did not detect any adaptations in C-bouton complexes. Firstly, certain motoneuron properties, such as C-bouton synapses, might be somewhat resistant to plasticity (Chalmers et al., 1991). Secondly, C-bouton synapses may already be organised to meet the increased demand following overload, and so even if it were more active, there may be no need for anatomical adaptation. And thirdly, C-bouton innervation may be sensitive to certain types of stimuli, but overload is either not appropriate or sufficent to stimulate plasticity.

In conclusion, our results show that a shift to a slower phenotype in EDL muscles is accompanied by a reduction in EDL motoneuron size, which perhaps allows EDL motor units to be recruited with less synaptic drive. The C-bouton complex, a key neuromodulatory synapse, is, however, anatomically unaffected by overload, suggesting that adaptation of these synapses was not neccessary. Whether C-bouton function adapts in other environmental conditions remains to be seen.

## Acknowledgements

We would like to thank Nadine Simons-Weidenmaier for technical support and Camille Lancelin for advice regarding confocal imaging and IMARIS analysis. The authors have applied a CC BY public copyright licence to any Author Accepted Manuscript version arising from this submission.

## Author contributions

R.W.P.K performed muscle experiments and analyses, spinal cord tissue preparation, interpreted results, wrote and edited the manuscript. A.G performed spinal cord experiments, acquired confocal images, analysed images using IMARIS, interpreted results, wrote and edited the manuscript. P.G.T performed muscle experiments and edited the manuscript. S.C & S.E provided technical advice and edited the manuscript. R.M.B helped design the study, interpreted the data, wrote and edited the manuscript. C.C.S conceptualised and designed the study, supervised experiments, analysed data, wrote and edited the manuscript.

